## Supplementary material 4 for "Improving yield and fruit quality traits in sweet passion fruit: evidence for genotype by environment interaction and cross-compatibility in selected genotypes"

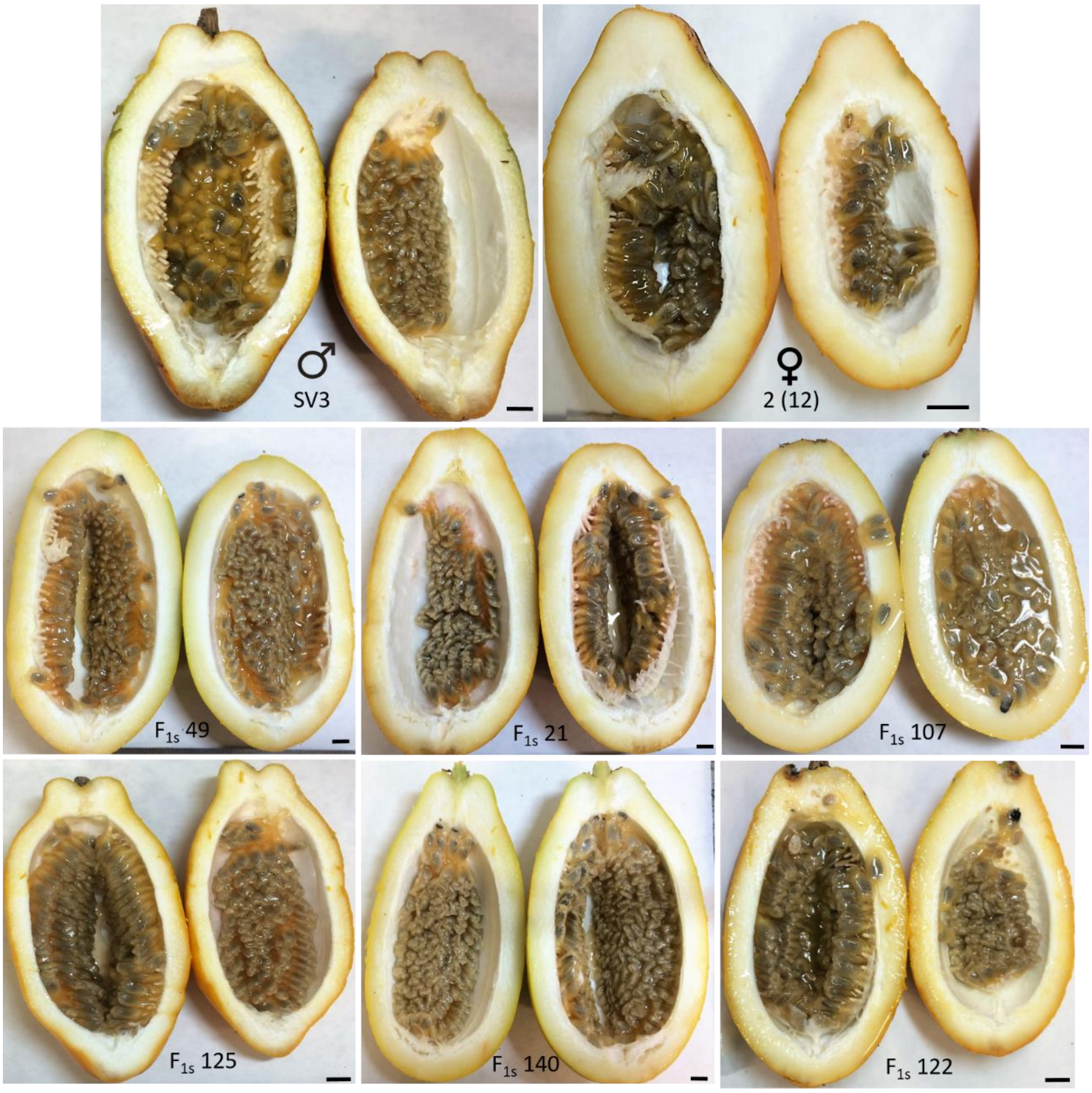


**Supplementary material 4**: Response to selection in sweet passion fruit. Note de difference in the thickness of skin and pulp content between fruits of the parental accessions (above). The six superior genotypes (below) were selected on the basis of a multiplicative index (MI). Bar = 1 cm.
