## Supplementary material 5 for "Improving yield and fruit quality traits in sweet passion fruit: evidence for genotype by environment interaction and cross-compatibility in selected genotypes"

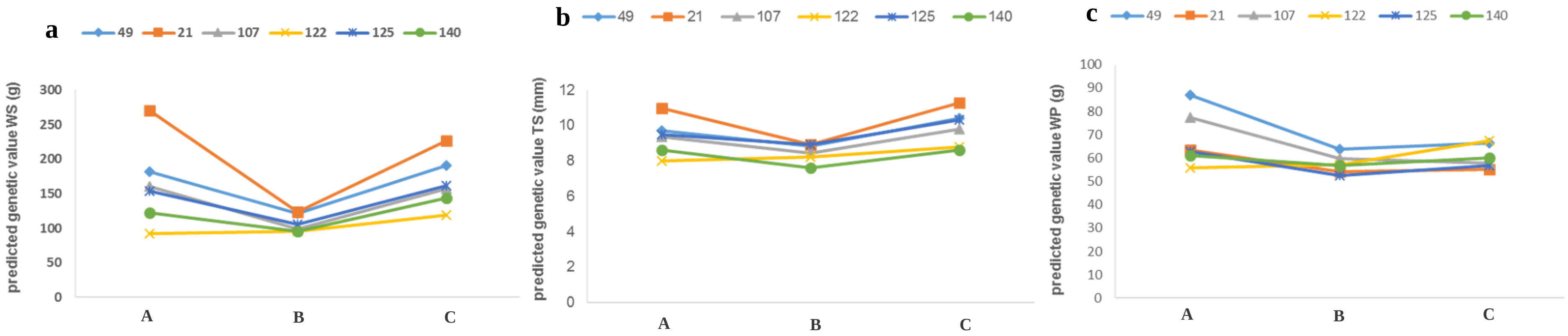


**Supplementary material 5**: Predicted genetic values of the six selected genotypes for Weight of Skin (a), Thickness of Skin (b) and Weight of Pulp (c) in each of the environments: A, B and C.
